# A Multiscale Model of Traumatic Brain Injury Suggests Possible Mechanisms of Post-Traumatic Psychosis

**DOI:** 10.1101/2021.04.19.440551

**Authors:** Dalton A R Sakthivadivel

## Abstract

Traumatic brain injury is a devastating injury to the brain that can have permanent or fatal effects, leading to life-long deficits or death. Among these effects is psychosis and schizophrenia, sometimes reported in the population of TBI sufferers. Here we evaluate a possible mechanism of post-traumatic psychosis, shedding light on the anomalous nature of psychosis as over-activity and brain injury as destruction. Using a multiscale model of the brain to relate molecular pathology to connectomic and macroscopic features of the brain, we identify cell lysis and membrane deformation as a possible mechanism for psychosis after injury. We also evaluate the reorganisation of functional networks and cortical activation post-injury, and find the features of a simulated brain under traumatic injury correlate with recorded results on the schizophrenic functional connectome. This provides a possible mechanism for post-traumatic psychosis, as well as a proof-of-principle of advanced multiscale modelling methods in computational psychiatry and neuromedicine. It also elaborates on the relationship between structure and function in the brain, information processing, and the delicate regulation of activity in healthy brains.

## Introduction

Traumatic brain injury (TBI), a result of a force sustained by the brain [1], is a disease with devastating effects on brain health, often resulting in loss of motor or cognitive ability, persistent vegetative state, or death [2]. Given the effects of both primary and secondary injury, TBI has chemical, physical, and mechanical implications, wherein the foundations of all neurological processes can be damaged [3–6]. Among the most destructive complications of TBI is axon injury, in which the physical connections of the brain are damaged. Axon injury describes tissue damage in the myelinated white matter tracts of the brain, caused by physical tissue deformation due to the force applied to the brain, or cytotoxic axon death immediately post-injury [7]. Such physical injuries, typically presenting as lesionation, are widely regarded as an instance of structural damage to the brain as opposed to a chemical process, due to the damage to brain architecture [8]. Axon injury can be focal or diffuse, describing the amount of localisation of axon damage [9–11], which is sometimes widespread, or can be localised to the site of the injury.

The aforementioned white matter tracts of the brain are often referred to as the brain’s structural connectome [12]. The necessity for physical structural connections is inherent to the nature of the brain, due to its organisation into cortical regions with independent generalised functions. These regions must be physically integrated to allow information to flow, which is accomplished by the structural connectome, where axon bodies serve as a medium for information travel. In axon injury, these damaged axons often experience decreased axonal transport due to degeneration of cellular structures [13–15]. Thus, when axon integrity and therefore transport in the structural connectome is disrupted by TBI, the brain’s essential communication network is expected to fail or exhibit decreased capability. Another way to characterise communication in the brain is by the abstract rather than the physical; that is to say, information flow itself, rather than the actual connections that may be utilised. The functional connectome, a counterpart to the structural connectome representing information flow, is a result of integration of brain regions, described by statistical correlations in spatio-temporal brain activity [16, 17]. Correlated activity in brain regions associates the regions as activated together and therefore processing the same signal, which is flowing between the two regions. Functional connectivity thus can indicate the amount of communication, or magnitude of information flow, between different brain regions. Resultantly, strength of connectivity indicates more information flowing between two regions, while functional connectivity networks represent the qualities of brain-wide cortical region activation. A popular viewpoint in connectomics research posits that aberrancy of functional connectivity is a measurable quantification of brain order and disorder. Accordingly, dysfunction of these brain networks may give rise to, or indeed reflect, larger disorder in the brain. In particular, we note that if functional networks represent the flow and subsequent processing of information across the brain, the normality of functional networks corresponds to the ability of the brain to process information in a regulated, ordered way.

While downregulation of information processing due to brain damage is expected, as suggested previously, there is a small but non-negligible portion of TBI patients that go on to develop either transient post-injury psychosis or chronic, long-term post-traumatic psychosis [18]. This is anomalous based on the damage expected to information processing networks in the brain, with psychosis typically being described as over-activity in key cortical networks [19, 20]. A 1969 review showed the incidence of post-TBI psychosis to be much higher than the average population, with further estimates of 0.9% to 8.5% increase in risk [21]. Others estimate a 60% increase in the risk of schizophrenia post-TBI [22]. It has been suggested that connectivity disorder created by trauma to the brain can be blamed for such disorders, by examining the specific correlation between lesionation and schizophrenia in the population [23]. More mechanistically, neuroimaging of a small number of patients with schizophrenic symptoms after cerebral injury has been shown in [24], which heavily imply a role of frontal or temporal lobe abnormalities in schizophrenic symptoms. In addition to structural pathology, there are chemical and cellular effects known to underpin schizophrenia, such as the abnormal neurotransmission of glutamate [25]. In TBI, parallely, there is the massive release of glutamate from dying cells, known to happen due to the lysis of cell membranes under shearing forces [26]. However—this has never explicitly been used to explain post-TBI psychosis. Clearly, while the increased relative risk of psychotic symptoms in TBI patients is apparent, and some tenuous connections have been drawn to facets of TBI and the pathophysiology or neurobiology of schizophrenia, the precise mechanism remains unclear. Uncovering such a molecular or cellular mechanism would mean a new avenue of attack to improve the cognitive recovery of all TBI patients, as well as those with post-traumatic psychosis specifically. It may also shed light on the delicate balance between excitation and inhibition in the brain, and what can come about when this balance is disrupted or dysregulated.

The aim of this study was to examine the possibility that TBI to the frontal lobe, and subsequent axonal lesions, could lead to an information processing disorder like schizophrenia, due to the structural disruption to axon bodies resulting in aberrant functional connectivity. In order to investigate possible mechanisms of post-traumatic psychosis, we require a comprehensive virtual model of the brain, due to constraints on what can be investigated in human subjects and the integration of various data modalities. In particular, integrating molecular information with behaviour in a mechanistic way is difficult, due to both the inaccessibility of such molecular measurements in the sensitive brains of TBI patients, and the different scale and resolution of molecular or cellular information as compared to connectomics and imaging data. Thus, we use a multiscale virtual model to simulate TBI and investigate the molecular correlates of possible psychotic symptoms. We do not merely take a computational approach to the problem, but apply biologically realistic virtual models to simulate brain activity and the effects of TBI. With a multiscale model, it is possible to investigate the problem faithfully at all levels of resolution, from cell bodies to brain cortices. Moreover, it is possible to evaluate functional connectivity in terms of actual information flow instead of approximations based on activity correlations.

As such, we use a multiscale model of the dynamical, hierarchical structure of the human brain, called The Virtual Brain (TVB). TVB employs a hierarchy of network-based scales, similar in structure to the human cortex, to model realistic brain corticodynamics [27]. At the lowest level it uses neural mass models, identical to physiologically observed cortical columns, and follows through to brain regions at the resolution of functional neuroimaging. A microscopic dynamical system for these neural populations is assigned to an array of connected spatial nodes in a so-called *brain network model,* which produces collective dynamics similar to those of macroscopic measurements like fMRI [28]. Thus, this biophysically realistic multiscale model combines parameter driven models of neural dynamics with observed macroscopic phenomena. We build a simulation of traumatic brain injury by focusing on structural aspects—lesionation to frontal lobe regions, represented by damaging network nodes—as well as dynamical aspects of TBI—modelling the effect of various diffuse molecular and cellular pathologies on the travel of electrical signals across the brain. This simulation is elaborated on in the Methods section. In particular, we evaluate the hypothesis that a combination of focal frontal lobe damage, and cell lysis leading to mass glutamate ejection, will lead to overactivity of the kind seen in psychosis, by having a direct effect on the functional network architecture and information processing capability of the brain. In so doing, we are able to verify whether or not these specific factors of TBI are connected to the signatures of schizophrenic-like symptoms and post-TBI psychosis that is currently poorly explained by the literature. In this study it was indeed found that patterns of functional connectivity are significantly different among TBI and healthy models, and that these differences correlate with the differences between brains of healthy and schizophrenic patients, as reported in literature on the subject. Thus, this study suggests how TBI can lead to schizophrenia, and also implies frontal lobe structural connectivity can influence aspects of functional connectivity.

## Results

20 simulations, each generated to have both EEG and BOLD fMRI results, were run with a ‘control’ or healthy design, using default parameter values and standard undamaged connectivity; 20 more were simulated with the specified TBI design. To avoid bias by simulation design, small amounts of noise were added to these waveforms, as well as variance in the parameter values themselves. Unpaired t-tests were run on the grouped average for each design, against the control. Simulation results focus on the ten regions typically associated with the Default Mode Network (DMN): the left and right dorsolateral prefrontal cortex (L/R DLPFC), the left and right temporal lobes (L/R TL), left and right parietal cortices (L/R PC), left and right components of the medial prefrontal cortex (L/R MPFC), and left and right components of the posterior cingulate cortex (L/R PCC).

### Default Mode Network Hyperconnectivity

Functional connectivity information was computed as a matrix of correlation strengths of simulated data, with mean connectivity matrices for each group in Figure 1. The difference between the functional connection between two regions, as compared between the healthy and TBI model, was found to be significant in 7 out of the 55 possible connections, with *p* < 0.05 (Table 1). Results tended towards stronger connectivity in the experimental group.

**Table 1.**
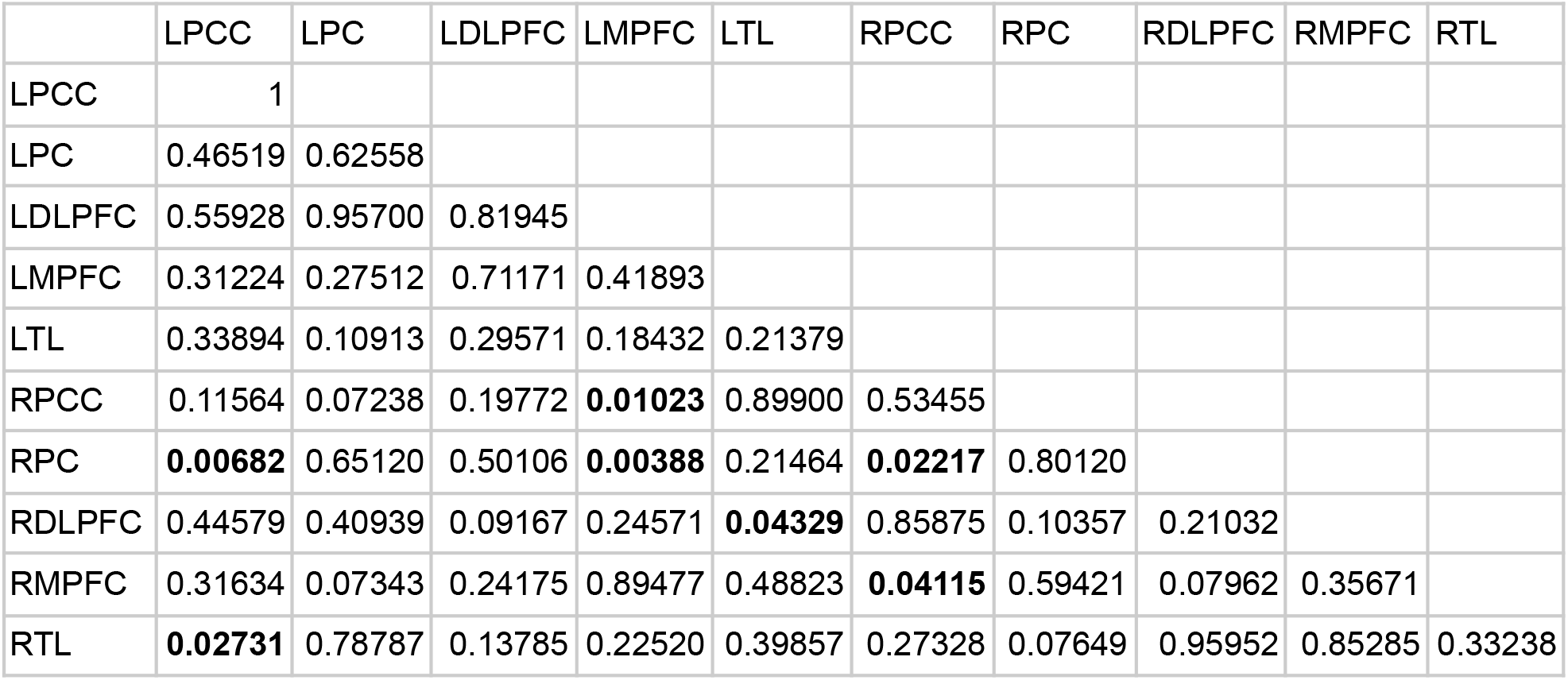
*p*-values for region-wise functional connectivity. The seven bolded entries denote significant and highly significant differences in connectivity at *α* = 0.05. Some marginally significant, but insufficiently low, p-values in the regime of *p* = 0.07 also appear.

|  | LPCC | LPC | LDLPFC | LMPFC | LTL | RPCC | RPC | RDLPFC | RMPFC | RTL |
| --- | --- | --- | --- | --- | --- | --- | --- | --- | --- | --- |
| LPCC | 1 |  |  |  |  |  |  |  |  |  |
| LPC | 0.46519 | 0.62558 |  |  |  |  |  |  |  |  |
| LDLPFC | 0.55928 | 0.95700 | 0.81945 |  |  |  |  |  |  |  |
| LMPFC | 0.31224 | 0.27512 | 0.71171 | 0.41893 |  |  |  |  |  |  |
| LTL | 0.33894 | 0.10913 | 0.29571 | 0.18432 | 0.21379 |  |  |  |  |  |
| RPCC | 0.11564 | 0.07238 | 0.19772 | **0.01023** | 0.89900 | 0.53455 |  |  |  |  |
| RPC | **0.00682** | 0.65120 | 0.50106 | **0.00388** | 0.21464 | **0.02217** | 0.80120 |  |  |  |
| RDLPFC | 0.44579 | 0.40939 | 0.09167 | 0.24571 | **0.04329** | 0.85875 | 0.10357 | 0.21032 |  |  |
| RMPFC | 0.31634 | 0.07343 | 0.24175 | 0.89477 | 0.48823 | **0.04115** | 0.59421 | 0.07962 | 0.35671 |  |
| RTL | **0.02731** | 0.78787 | 0.13785 | 0.22520 | 0.39857 | 0.27328 | 0.07649 | 0.95952 | 0.85285 | 0.33238 |

**Figure 1.**
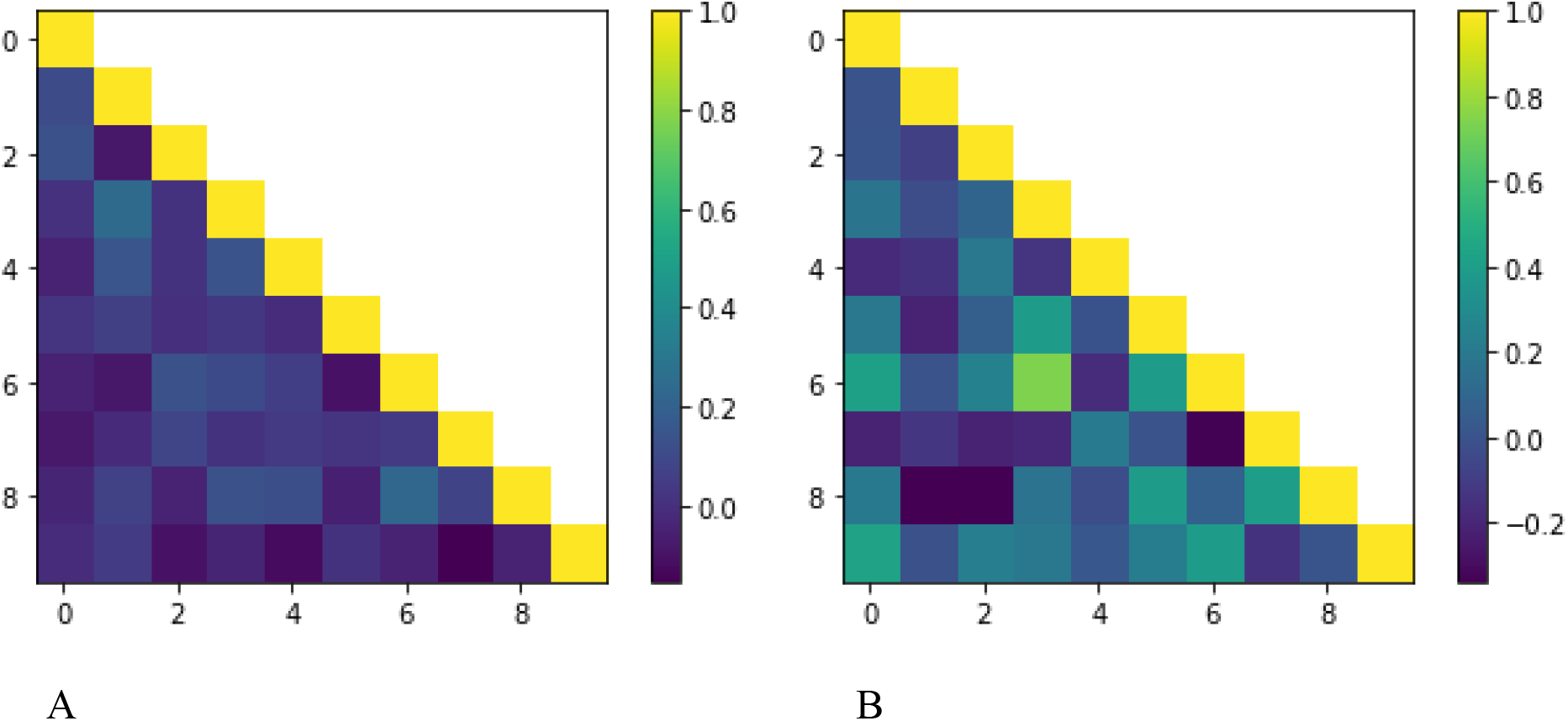
Hyperconnectivity in mean region-wise correlation. Figure 1A depicts the correlation matrix for the control group, and Figure 1B, the experimental. The experimental group clearly exhibits higher mean functional connectivity, with significance tests in Table 1.

In the healthy model, the strongest off-diagonal connection is between the LPC and LMPFC, with an average strength of 0.2473. This is not the case in the TBI model, with the strongest connection running between the LMPFC and the RPC, having an average strength of 0.7408. Here, the healthy model had only a weak connection (0.1039). The difference between the LMPFC-RPC connections in the two models is significant at *p* < 0.005. Another significantly different connection existed between the LMPFC, and the RPCC, having a strength of 0.4002 (experimental) and 0.03467 (control). We also have significant hyperconnectivity between the LTL and the RDLPFC, with *p* < 0.05.

As instances of PCC hyperconnectivity, both left and right components of the PCC had significantly stronger connections in the experimental model, with LPCC-RPC, LPCC-RTL, RPCC-RPC, and RPCC-RMPFC all being significantly different. The respective connectivities are given in Table 2.

**Table 2.** Elevated connectivity in the Posterior Cingulate Cortex. Table shows the connectivity values for each group’s PCC, where the difference was significant. Hyperconnectivity in the PCC is observed in the experimental group, where the correlation is significantly higher than control.

|  | LPCC-RPC | LPCC-RTL | RPCC-RPC | RPCC-RMPFC |
| --- | --- | --- | --- | --- |
| Control | -0.043407 | -0.005746 | -0.097472 | -0.050174 |
| Experimental | 0.421132 | 0.432929 | 0.402546 | 0.402101 |

### Network Shape and Reorganisation

To compare the presence or absence of network links, a threshold was applied to remove links that were close to zero and had insignificant correlations. In comparing the two networks this way, multiple connections were not common to both models, suggesting network reorganisation (Figure 2). The thresholded network in the healthy model is relatively equally distributed among and across hemispheres, with arguably fewer interhemispheric connections when noting the relative differences in connective ‘capacity’ (more possible interhemispheric connections, such that the percentage unconnected is higher). The TBI model is weighted towards interhemispheric connections both comparatively and in percentage connected, with many connections in the lower left quadrant of the matrix, between right-lying and left-lying regions. Many such connections are absent in the healthy model.

**Figure 2.**
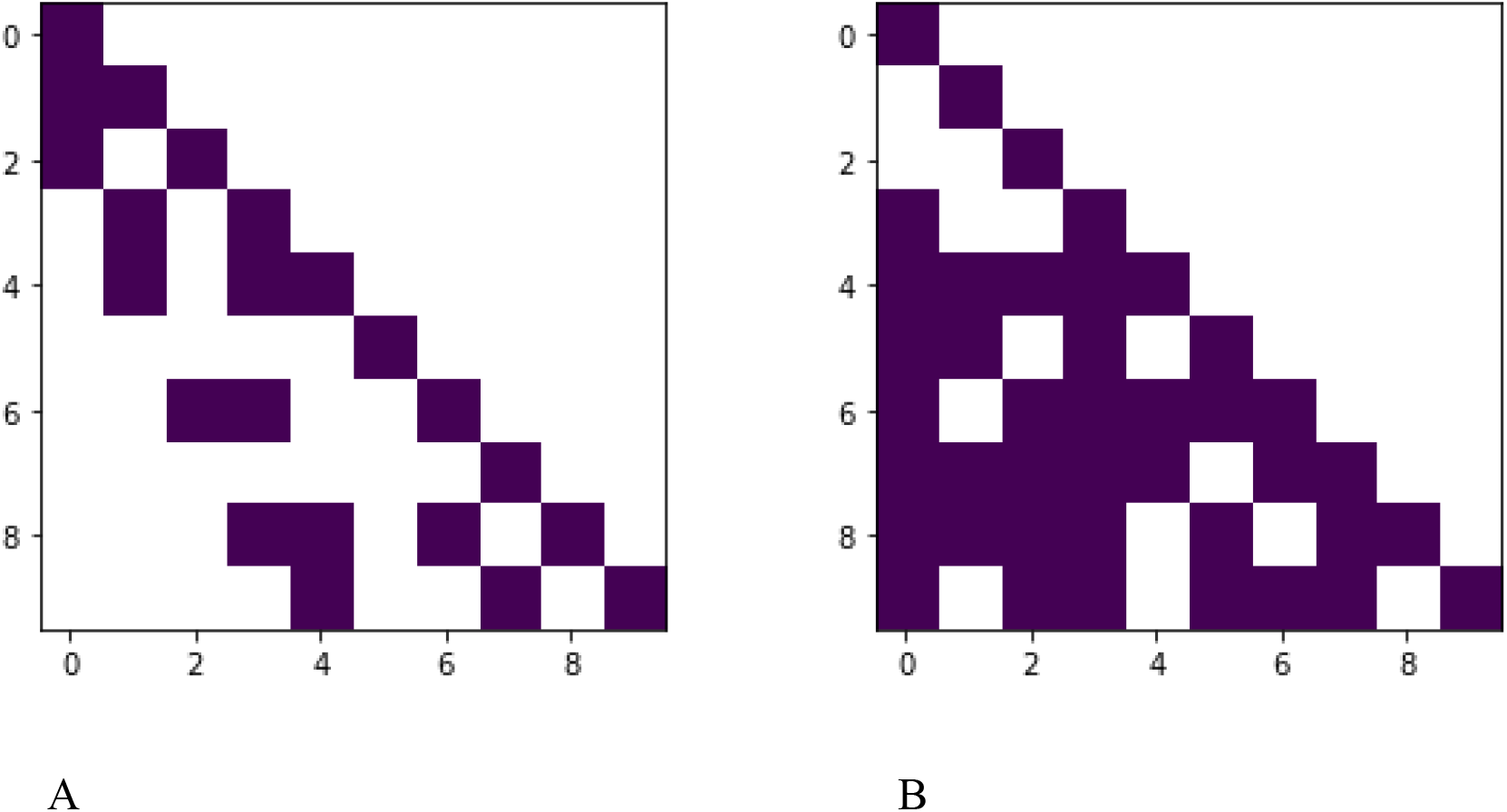
Binarised, thresholded network shows many more connections in the experimental group than in the control. Figure 1A depicts the binarised correlation matrix for the control group, and Figure 1B, the experimental. An arbitrary threshold of *r^2^* > 0.01 was chosen to determine a ‘significant’ correlation, such that the magnitude of any correlation must have exceeded 0.1 to be counted. When this threshold was applied, the network structure showed reorganisation in the experimental group, having many more significant connections than the control.

Graph theory analyses were performed on both networks to characterise the shape of each. With respect to quantification of neurological phenomenon, three measures were found useful to record: global clustering coefficient (*C*), average degree (*K*), and neighbourhood degree, or average number of edges in any connected subgraph (*N*). In the healthy model, both *C* and *K* were lower at 0.3750 and 2.2453 respectively, where in the TBI model they were 0.6467, and 2.5950, respectively. This indicates a higher tendency to form sub-networks, or clusters, and for those clusters to be highly connected.. *N* tended to higher values in the experimental model, confirming this tendency to form dense clusters with high degree nodes being more likely to connect to other high degree nodes, in what is called preferential attachment. *K* being greater in the TBI model than in the healthy model, indicates denser connections being formed.

### Cortical Hyperactivity in DMN Regions

Also telling are graphs of cortical region activation. The underlying quality which gives rise to functional connectivity is cortical activation, because functional connectivity is a measure of the reception and inflow of information to a specific region from another. In the TBI model, elevated cortical source activation in the LTL (Figure 3A and 3D) and RDLPFC (Figure 3C and 3F) areas correspond to hyperconnectivity of these regions. In our dataset, four out of the eight condensed DMN regions exhibited highly significant overactivation in the TBI model (p < 0.001). The indicated overactivation was found in the LTL, RDLPFC, LLPC and RLPC.

**Figure 3.**
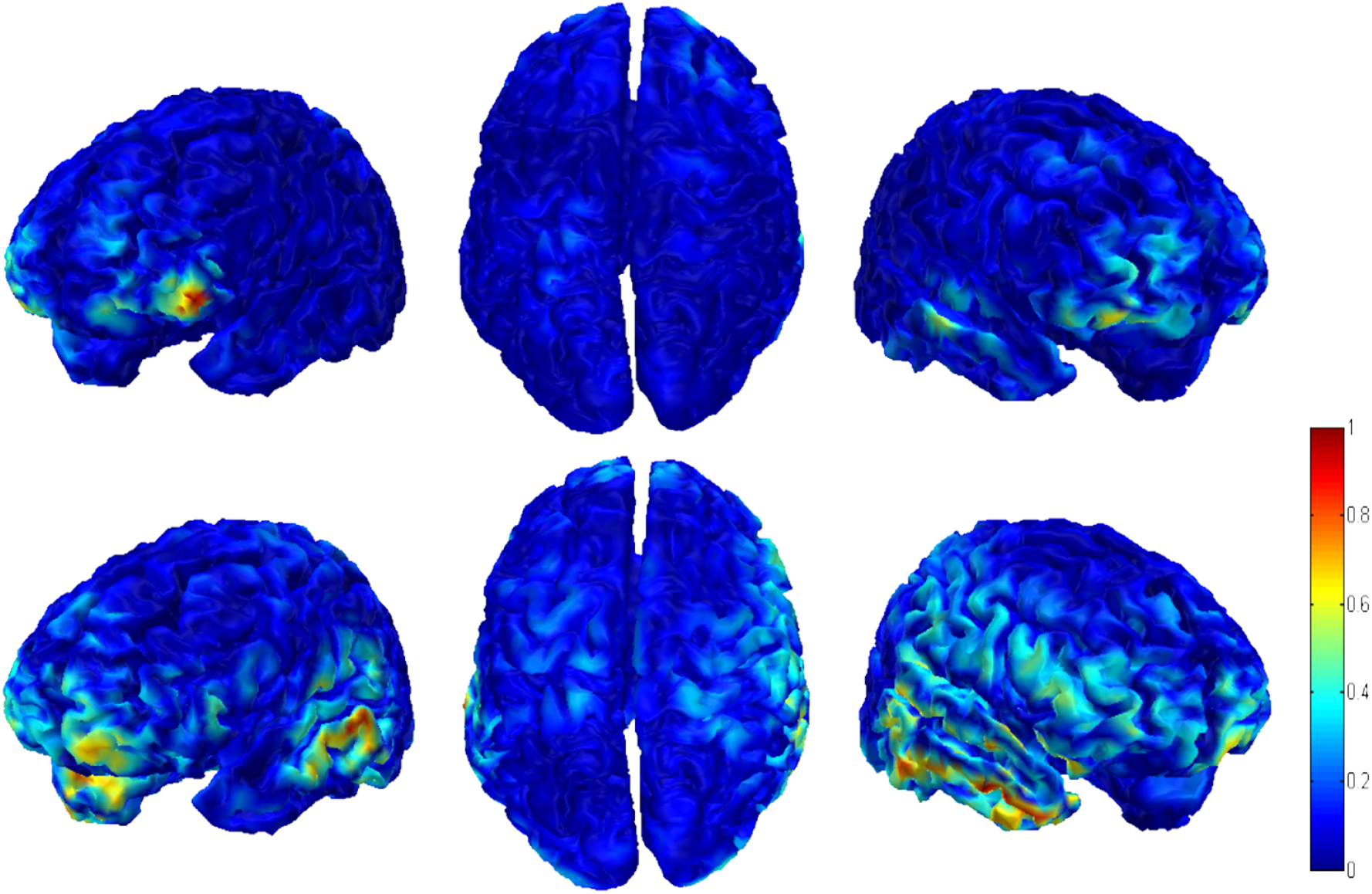
Cortical overactivation present in DMN regions in TBI simulation. Top row, 3A-C, shows mean cortical current density calculated on a healthy simulation, while the bottom row, 3D-F, shows the same result for TBI simulations. A larger radius of activity and more intense activity level is observed in the TBI model.

## Discussion

As compared to the healthy model, in the TBI model, the brain exhibited both weakened and strengthened portions of resting state connectivity. Of interest was the difference in magnitude of incoming and outgoing connections to the RDLPFC, as literature correlates overactivation and hyperconnectivity of the RDLPFC with schizophrenic symptoms [29–31]. This is perhaps due to the DLPFC being responsible for efficient processing of cognitive information [32]. Deviations in this area would conceivably lead to symptoms of schizophrenia like hallucination or paranoia, where cognitive information being processed does not match that of reality. Also of interest was the behaviour of the LTL. Hyperconnection of the LTL is correlated with schizophrenic symptoms [33, 34]. A probable theory is that due to the position of the LTL as a processing element in the ventral stream, responsible for processing visual and auditory stimuli, higher than normal information flow could lead to generation of non-existent stimuli, such as what is seen in auditory and visual hallucinations. Further, both left and right lateral parietal cortices exhibited overactivation in the TBI model, though they did not exhibit hyperconnectivity. Aberrations in the parietal cortex are tentatively linked to schizophrenia [35], with a possible explanation for this link being the responsibility of the parietal cortex in processing spatial information and perception [36]. When gone awry, this may complicate the distinction between what is real in space and what is not.

An additional piece of analysis involved application of graph theoretical measures to the functional connectomes of both models. It is widely reported that a particular organisation of networks, termed the small world network (SWN), is optimal for healthy and effective brain communication. When this organisational scheme is departed from, it often can result in schizophrenia [37–40]. In particular, the definition of an SWN is a network with a high tendency to cluster around given nodes, making small sub-communities, combined with a low average distance between nodes [41]. This makes for efficient communication and dense connections between nodes. The TBI network was far more efficient than the healthy brain, with denser connections and more hubs of communication. This once again relates to the efficiency of communication in the network: hubs serve to integrate different portions of the network and are necessary for the sustained ease of communication in the network. It is possible that strong hubs forming could lead to over-processing of information, in such a way that false information or experience is generated. The usefulness of graph theoretical analysis is limited by the size of the data set; usually these measures are applied to much larger brain networks, unlike the DMN, and are subject to statistical analysis based on a high population size. This study’s application of graph theoretical measures are limited by its small sample size of one network per simulation and the small network itself.

The significance of this study is twofold: for one, it addresses portions of the controversy over the effect of structural connectivity on functional connectivity, implying that aberrancy in frontal lobe structural connectivity can change aspects of functional connectivity in the frontal lobe and other regions of the brain. It also suggests that the effects of TBI can put a patient at risk for developing schizophrenic symptoms commonly associated with overactivity or hyperconnectivity of particular brain regions. This is an as yet unexplored possibility, with postulated but not proven links between the two diseases. While it is accepted that due to the delicate nature of the brain, one form of disorder can lead to other forms of disorder in the brain, with dementing disorders or disorders like post-concussive syndrome pointed to as a consequence of TBI, schizophrenia is a complication only touched upon and never before quantitatively investigated.

A practical limitation of this study is that it implies immediate onset of schizophrenia upon focal lesion presentation. What is perhaps a more likely interpretation is that TBI and resultant axon injury may lead to onset of schizophrenic-like symptoms, such as hallucinations or psychosis, and perhaps future persistence of these symptoms. Considering the function of the DMN, activated during resting state activity, presentation of these symptoms would make sense; as a centre for self-reflection, overactivity of portions of this network may indeed result in less distinction between what is inner and what is outer. In the case of individuals with increased activation or connection of DMN regions, the DMN may not behave in a normal, stimulus load dependent manner, instead continuing to function alongside stimulus processing [19]. It is possible that this would lead to inner ideas and self-constructed ‘stimuli’ being perceived as existent in reality, in a fashion consistent with the definition of a hallucination, paranoia, or psychosis. Individual regions may also play a role in this: considering the function of the DLPFC within the dorsal stream and thus its responsibility in methods of processing of external stimulus, overactivity may lead to over processing of reality. Stimuli may be constructed in such a way that there is a deviation from the way a given stimulus is actually present and the way the brain and particularly the DLPFC interprets it to be so. This could conceivably lead to a hallucination or paranoid episode. Along these lines, the increase in connection strength into the PCC in the TBI model is notable. Deviations in the PCC are correlated with schizophrenia in literature [42–44]. The PCC, which is both part of the dorsal and ventral streams and mediates large portions of the DMN, may be implicated in both mechanisms of processing dysfunction. In fact, the PCC is of particular interest due to its nature as a central hub of information processing. Given this function, aberrancy in this area of the brain may help to unify the ideas of disorder in information flow from structural damage and disorder in information processing. When this area is damaged structurally, the effects on stimulus and information processing would be wide reaching.

The suggested trend of structural damage leading to increased connectivity challenges the perhaps more intuitive idea of TBI, which initially might lead to one saying connectivity would be downregulated due to physical destruction. It is reasonable, however, considering that the brain may process information in an erratic way due to the relationship between healthy and abnormal information conveyance in the brain. The physical brain is intimately linked to the 'abstract’ brain. In other words, the structure of the brain is linked with cognitive processes, and disruption of the brain can result in disruption of cognition, corroborating known results in TBI [45].

## Methods

We utilised a multiscale model of the human cortex, The Virtual Brain [27], to investigate the effect of chemical imbalances on brain network organisation. The brain, being organised in a multiscale and hierarchical fashion, encodes information about genes and neurotransmitters, single neural cells, and local networks of neurones in its macroscopic behaviour, such that fMRI is a measure of collective activity, in addition to its own dynamical process. Thus, connecting these scales is valuable for the problem posed here, and a multiscale model was used.

The structure of a simulation in TVB consists of two primary data: local and global connectivity, defining the topologies of the networks on which things are simulated, and the neural model that is being simulated. Beginning with the lowest scale, TVB uses neural mass models (NMMs), mean field models of the activity of multiple individual neurones in tightly coupled populations. Mean field theory is a method from statistical physics that models the collective activity of a group of objects, if they have well-defined ‘mean’ dynamics that can summarise the population well [46]. These neural populations thus encode information about the firing dynamics of single neurones, as well as any synaptic or chemical information encoded in a given model. These neural mass models are coupled together into a local network, described by a larger equation in terms of the NMM and the couplings or interactions between close-lying masses. The network itself, made of systems of equations of multiple NMMs, are assembled into a matrix in such a way that the nodes of a local network are composed of *m* neural masses. Then, *j* local networks are assembled into a global network, where networks of local networks define regions subject to structural connectivity interactions, the conduction velocity mediating the interactions on these nodes, and time-delays between signals. The coupling topology at each level is an adjacency matrix between equations, and the interactions in this matrix mediate the dynamics of that level as well as the levels above it, by virtue of coupling. The precise mathematical formulation of this brain network model is found in [28]. Single node dynamics were calculated in TVB using an extension of the Larter Breakspear model,

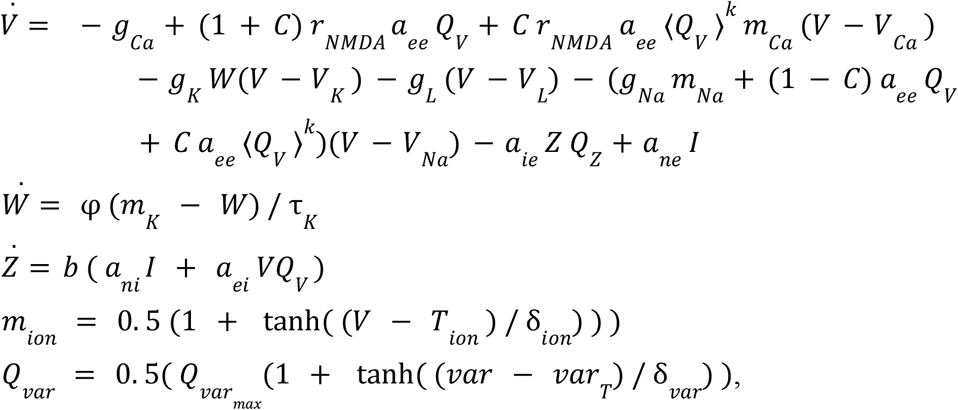

with *k* being a node in the network. Default values and parameter definitions can be found in^1^. Ion values can be Ca, Na, K; variables can be *V, W,* and *Z*.

We determined two important features of the simulation to be connectomic lesionation and synapto-chemical neural mass model parameters. We simulated left hemispheric frontal lobe lesionation by decreasing the weights of outgoing connections from three key subregions of the left posterior frontal cortex: the LPFCPOL, LPFCVL, and LPFCCL. These regions are not only highly connected and located in the frontal lobe, the seat of information rocessing, but are vulnerable to focal injury due to their forward facing positions, close to the forehead. Outgoing connections from these regions were chosen randomly for deletion, modelling the unpredictability of cell structure integrity. The particular connections that were destroyed were chosen to not be DMN regions, assuming that if the exact cognitive effects found are shown to be independent of the roles of these regions, we can reasonably assume there is some more fundamental network reorganisation and the results shown extrapolate to other lesion topologies.

Next, we simulated the neurochemical pathology of TBI. It has been shown that non-linearities in mean field neural dynamics have identifiable signatures of chemical or ion dynamics [47, 48], so, we sought to identify a neural mass model capable of modelling this. As shown above, we utilised the Larter-Breakspear model of ionic kinetics in neural firing to model how mean field neural firing dynamics depends on parameters of the synapse and membrane [49, 50]. We based our parameter manipulations off of chemical pathologies recorded in [3, 5, 6, 15, 51], with estimations of parameter values taken from [52, 53]. Specifically, membrane conductances were increased to model the effect of TBI on cell membrane integrity, and ionic Nernst potentials were increased to correspond to the increase in extracellular concentration of all ions. A stable regime of parameters that gave dynamical oscillations was identified using the nullclines of the phase plane of the system, so as to simulate living but unhealthy brain tissue, similar to an awake and recovering TBI patient.

Raw neural time-series were recorded as the readout of region-wise ensemble averaged activity. EEG and fMRI were simulated by lead-field matrix transformation and HRF kernel convolution, respectively, generating realistic waveforms for both data modalities. Cortical current density for ten DMN regions was estimated in eConnectome [54] and functional connectivity was calculated using region-wise Pearson correlation on the fMRI signals. Graph theory analyses were performed in Python3 using the NetworkX package [55].

## Conclusion

In this paper, a multiscale model was used to show that a particular structural and neurochemical pathology found in TBI can lead to signatures of schizophrenia or psychosis in the brain. Neural phenomena associated with TBI were simulated, especially excitatory cell lysis and the resultant disruption to ion dynamics. Signatures of psychosis, such as hyperconnectivity in the DMN, network reorganisation and hub formation, and overactivation, were confirmed as results of intentional parameter manipulations at the lowest level of the model. A unique model in its ability to act as a ‘sandbox’ for the brain, such a multiscale model allowed the various relationships between pharmacological pathology and observed behaviour to be examined in an end-to-end fashion. More broadly, such models aim to answer a question definitively, through testing a hypothesis by simulation; in this case, can cell damage and resultant aberrations in neurochemical ion kinetics result in the psychosis that is sometimes observed post-TBI? These simulations suggest this is the case, pointing us in the direction of pharmaceutical treatments for this disorder, a better understanding of the brain, and a use-case for multiscale modelling methods in neuropsychiatry.

1 http://docs.thevirtualbrain.org/api/tvb.simulator.models.html#module-tvb.simulator.models.larter_breakspear

## Notes

### Competing Interest Statement

The authors have declared no competing interest.

